# Deterioration of dry skin in arthritis model mice via stress-induced changes in immune cells in the thymus and spleen

**DOI:** 10.1101/641720

**Authors:** Kenji Goto, Keiichi Hiramoto, Ion Takada, Kazuya Ooi

**Affiliations:** Laboratory of Clinical Pharmacology, Faculty of Pharmaceutical Sciences, Suzuka University of Medical Science, Suzuka, Japan; Laboratory of Pathophysiology and Pharmacotherapy, Faculty of Pharmaceutical Sciences, Suzuka University of Medical Science, Suzuka, Japan

**Keywords:** Arthritis, Dry skin, Stress, T helper 1 cell, T helper 2 cell, T helper 17 cell, regulatory T cell, Thymus, Spleen, Corticosterone, Interleukin-6

## Abstract

Skin dryness is a characteristic of rheumatoid arthritis model mice. We previously reported that the stress hormone glucocorticoid (i.e., corticosterone) is related to the induction of dry skin in arthritic mice. However, the mechanism through which stress induces dry skin in these mice is still unclear. Therefore, in this study, we examined the relationship between stress and induction of dry skin in arthritic mice. Physical stress load in mice with DBA/1JJmsSlc collagen-induced arthritis was treated with water immersion stress, and transepidermal water loss and the expression of markers associated with allergic reactions and inflammation was evaluated. Deterioration of skin dryness was observed in stressed arthritic mice compared with that in unstressed arthritic mice. Moreover, plasma levels of interleukin-6 and corticosterone were increased in stressed arthritic mice compared with those in unstressed arthritic mice. We also observed decreased regulatory T cell numbers and increased T helper type 2 cell numbers in the thymus of stressed arthritic mice compared with those in unstressed arthritic mice. These results suggested that abnormalities in the immune system were related to deterioration of dry skin in stressed arthritic mice. Thus, reduction of stress may prevent deterioration of dry skin in mice with arthritis.

## INTRODUCTION

Rheumatoid arthritis (RA) is an autoimmune disease characterized by progressive destruction of the synovial membrane and bone with persistent chronic inflammation as the main symptom [1, 2]. Nonarticular lesions, such as blood vessels, lungs, kidneys, and skin, are observed in approximately 40% of patients with RA [3]. Rheumatoid nodules and xerosis occur as skin lesions in RA [4, 5] and can affect patient quality of life [6]. Decreased skin barrier function (i.e., dry skin) causes itching [7], and injury to the stratum corneum caused by scratching can then induce skin infections [8, 9]. Therefore, elucidation of the mechanisms regulating the induction of dry skin and approaches for preventing dry skin is necessary to avoid such adverse reactions.

We previously reported that dry skin occurs in an arthritic mouse model via a mechanism involving mast cells and histamine secreted from mast cells [10]. In addition, reactive oxygen species (ROS), neutrophils, and thymic stromal lymphopoietin were found to induce dry skin via mast cells and expression of cytokines (e.g., interleukin [IL]-6 and tumor necrosis factor-α) [11]. Thus, ROS released from neutrophils stimulates thymic stromal lymphopoietin, and activated thymic stromal lymphopoietin induces mast cells, which degranulate histamine and release IL-6, resulting in dry skin.

Patients with RA also suffer from stress owing to severe joint pain. Psychological stress causes increased glucocorticoid (GC) production and stimulates hypothalamic corticotropin-releasing hormone production, leading to increased secretion of adrenocorticotropic hormone by the pituitary gland [12]. These changes caused increased synthesis and secretion of GC from the adrenal glands. GC has anti-inflammatory effects, but induces abnormalities in the structure and function of the skin by upregulating Toll-like receptor 2 [13]. Furthermore, treatment with the GC receptor (GCR) inhibitor RU486 ameliorates dry skin in arthritic mice [11]. Accordingly, stress is related to induction of dry skin in arthritic mice. Additionally, stress is involved in the onset of RA itself [14]. However, despite these studies, the mechanisms through which stress induces dry skin in arthritic mice are still unclear.

In this study, we evaluated the effects of water immersion stress on dry skin in arthritic mice.

## MATERIALS AND METHODS

### Animals

Specific pathogen-free DBA/1JJmsSlc mice (10 weeks old) were obtained from Japan SLC (Hamamatsu, Shizuoka, Japan). We used these mice as a collagen-induced arthritic mouse model. The method for inducing arthritis in these mice was described previously [11]. The mice were housed under a 12:12-h light/dark cycle at a constant temperature of 23°C ± 2°C and 55% ± 10% relative humidity. The mice had free access to water and laboratory chow (CE-2; Oriental Yeast Co., Tokyo, Japan). All animals were treated in accordance with the Guide for the Care and Use of Laboratory Animals of Suzuka University of Medical Science (approval number: 66). All experiments were performed under sodium pentobarbital anesthesia.

### Experimental design

After allowing the mice to acclimate for 1 week, stress treatment was applied for 4 days, and mice were sacrificed on day 5 (Fig. 1). On day 4, the hair on the dorsal skin was clipped, and a depilatory cream (Veet Hair Removal Cream Tube Fit Sensitive; Reckitt Benckiser, Slough, UK) was used to remove the remaining hair to allow evaluation of dry skin. We confirmed that the hair cream did not cause skin abnormalities in the mice. Skin, thymus, spleen, and blood samples were collected on the final day of the experiment. Untreated DBA/1JJmsSlc mice and DBA/1JJmsSlc collagen-induced arthritic mice served as the control and arthritis groups, respectively (n = 5 per group). The arthritic mice were randomly assigned (n = 5 per group) to the arthritis and arthritis plus stress groups.

**Figure 1.**
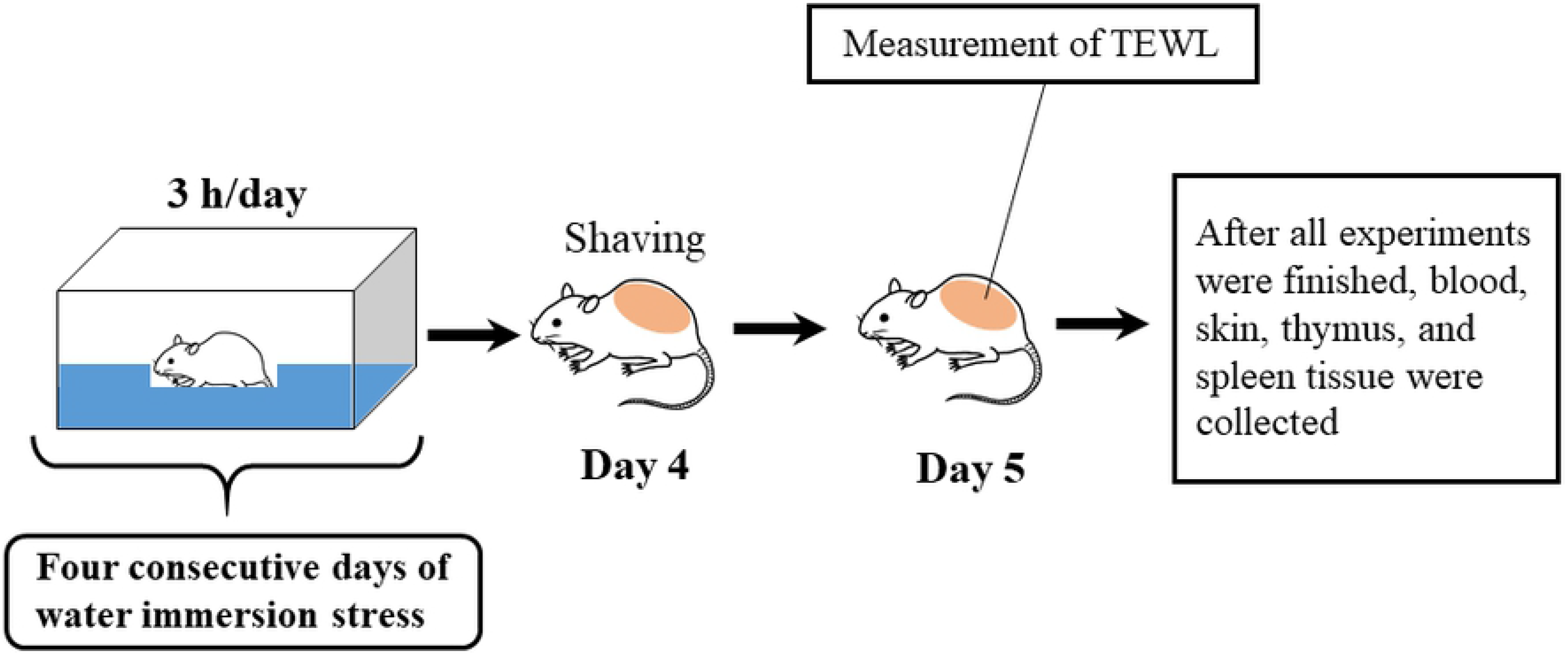
Schematic of the procedure for this experiment. TEWL: transepidermal water loss.

### Immobilized-water immersion stress

Water immersion stress was performed as described by Izumi et al. (1983) [15], with some modifications. Briefly, for water immersion stress, animals were confined to a plastic box (276 × 445 × 204 mm) and immersed in 300 mL water (25°C) for 3 h on days 1–4. The stress levels of the mice were confirmed by the increase in corticosterone levels.

### Measurement of TEWL

On the final day, TEWL in the dorsal skin of each mouse was measured. TEWL serves as a marker of skin permeability, reflecting the barrier function of the skin; increased TEWL indicates dry skin. Measurements were carried out using a Tewameter TM300 (Courage + Khazaka Electronic GmbH, Cologne, Germany) [16]. Values were recorded once the reading had stabilized, typically 10 s after the probe was placed on the skin. Data are presented as the average of three independent measurements.

### Quantification of IL-6, corticosterone, hyaluronan, and ROS levels

On the final day of the experiment, mice were anesthetized by intraperitoneal injection of sodium pentobarbital (50 mg/kg; Nacalai Tesque, Kyoto, Japan), and blood samples were collected by cardiac puncture. Plasma was separated from blood samples by centrifugation at 3000 × *g* for 10 min at 4°C, and the supernatant was used for analysis. Plasma levels of IL-6, corticosterone, and hyaluronan were measured using commercial enzyme-linked immunosorbent assay kits (IL-6: BioLegend, San Diego, CA, USA; corticosterone: Enzo Life Science Inc., Farmingdale, NY, USA; hyaluronan: R&D Systems, Minneapolis, MN, USA), according to the manufacturers’ instructions. ROS levels were determined using an OxiSelect in vitro ROS/RNS assay kit (STA-347; Cell Biolabs, Inc., San Diego, CA, USA), according to the manufacturer’s instructions. Optical density was measured with a microplate reader (Molecular Devices, Sunnyvale, CA, USA).

### Preparation and staining of dorsal skin

Mice were sacrificed 5 days after the start of the experiment. Dorsal skin tissue samples were isolated and fixed in phosphate-buffered saline containing 4% paraformaldehyde (Wako Pure Chemical Industries, Osaka, Japan). Fixed tissue specimens were embedded in Tissue Tek OCT Compound (Sakura Finetek, Tokyo, Japan) and frozen. The tissue blocks were cut into 5-μm-thick sections that were stained with H&E for histopathological analysis of the tissue according to conventional procedures. H&E stains the nucleus and cytoplasm, allowing histopathological determination of tissue pathology. We microscopically evaluated H&E-stained skin tissues. To determine overall skin thickness, 10 regions in which the skin appeared flat in acquired images were randomly selected. The length from the outer layer of the epidermis to the border of the subcutis was measured, and the average value was calculated. Additionally, skin specimens were stained with toluidine blue to visualize mast cells. We microscopically evaluated skin tissues stained with toluidine blue according to conventional procedures. Mast cells were quantified by counting cell numbers per mm^2^ field in 10 randomly selected regions. Furthermore, Masson’s trichrome staining was performed using a Trichrome Modified Masson’s Stain Kit (Scytek Labs), according to the manufacturer’s instructions. We microscopically observed collagen in the skin.

### Western blotting

Dorsal skin, thymus, and spleen samples were homogenized in lysis buffer (Kurabo, Osaka, Japan) and then centrifuged at 8000 × *g* for 10 min. The supernatant from each sample was collected and stored at −80°C until analysis. After thawing, equal amounts of protein (skin: 5 μg/lane, thymus: 9.75 μg/lane, spleen: 30.5 μg/lane) were loaded onto 4–12% Bis-Tris Bolt gels (Life Technologies, Carlsbad, CA, USA) and electrophoretically separated at 200 V for 20 min. The proteins were then transferred using an iBlot Western blotting system (Life Technologies) to nitrocellulose membranes. The membranes were blocked overnight at 4°C with 5% skim milk and then incubated at 25°C for 1 h with primary antibodies against GCR (1:1000; Bioss Antibodies, MA, USA), Annexin V (1:1000; Abcam, Cambridge, UK), T-bet (1:1000; Abcam), Foxp3 (1:1000; MBL, Aichi, Japan), RORγt (1:1000; Biorbyt, Cambridgeshire, UK), GATA3 (1:1000; Cell Signaling Technology, Danvers, MA, USA), and β-actin (1:5000; Sigma-Aldrich, St. Louis, MO, USA). The membranes were then incubated with horseradish peroxidase-conjugated secondary antibodies (Thermo Fisher Scientific, Waltham, MA, USA), and the signal was detected with ImmunoStar Zeta (Wako Pure Chemical Industries) and a lumino-image analyzer (LAS-4000; Fuji Film, Greenwood, SC, USA). Protein bands were analyzed by densitometry, and the signal intensity was normalized to that of β-actin.

### Statistical analysis

Data are presented as means ± standard deviations. Tukey’s post-hoc tests were used to evaluate differences between groups, and results with *P* values of less than 0.05 were considered statistically significant.

## RESULTS

### Changes in skin condition

A schematic of the experimental model is given in Fig. 1. First, we measured transepidermal water loss (TEWL) in the dorsal skin of mice to evaluate skin dryness (Fig. 2A). The results showed that arthritic mice had higher TEWL than control mice and that stressed arthritic mice had higher TEWL than arthritic mice. In addition, we examined histological changes in the skin of these mice using hematoxylin and eosin (H&E) and Trichrome staining (Fig. 2B and 2C). The skin of arthritic mice was thicker than that of control mice; however, the skin of stressed arthritic mice was thinner than that of arthritic mice and similar to that of control mice. Collagen levels were decreased in the skin of arthritic mice compared with that in control mice and decreased in the skin of stressed arthritic mice compared with that in arthritic mice.

**Figure 2.**
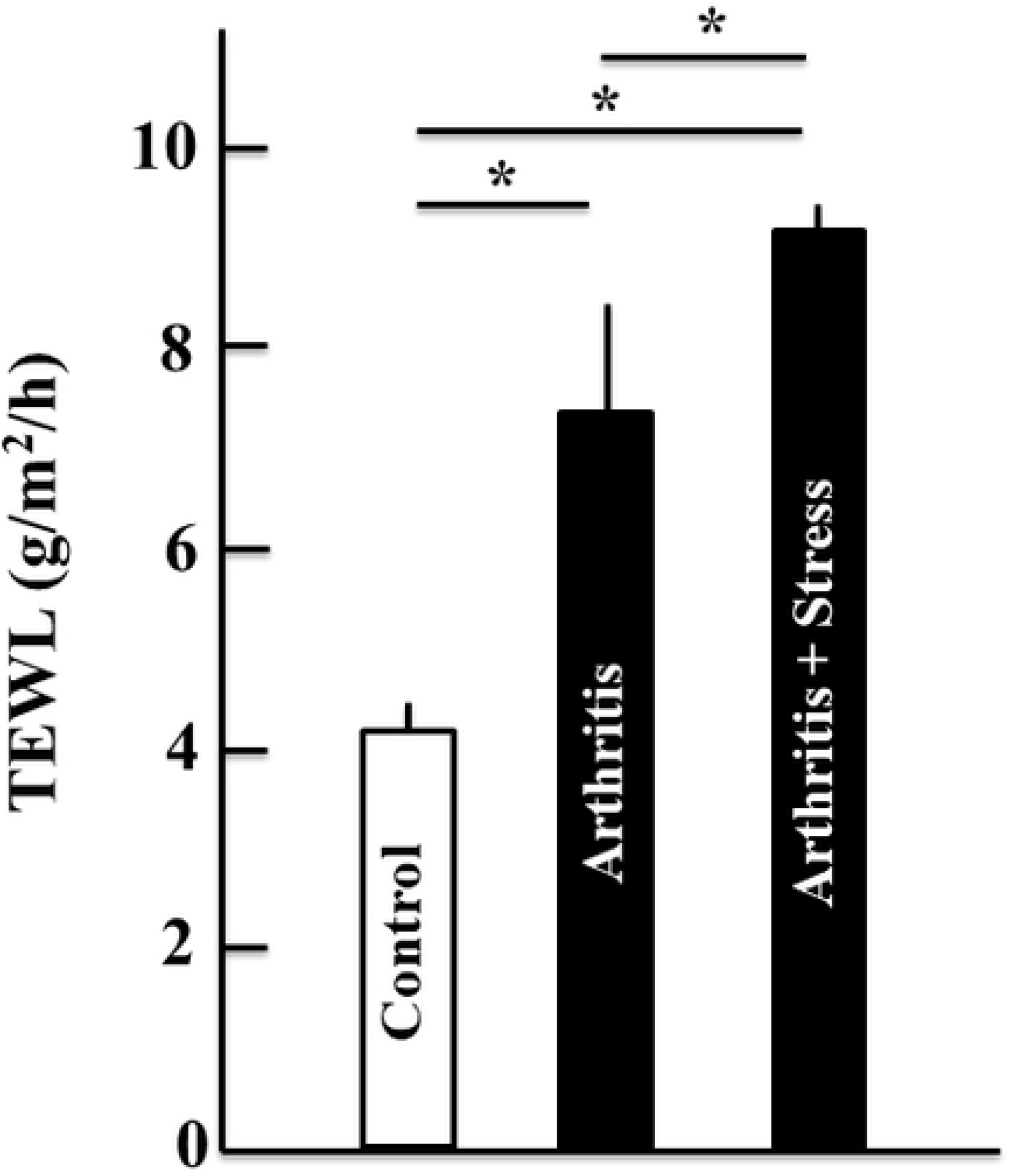
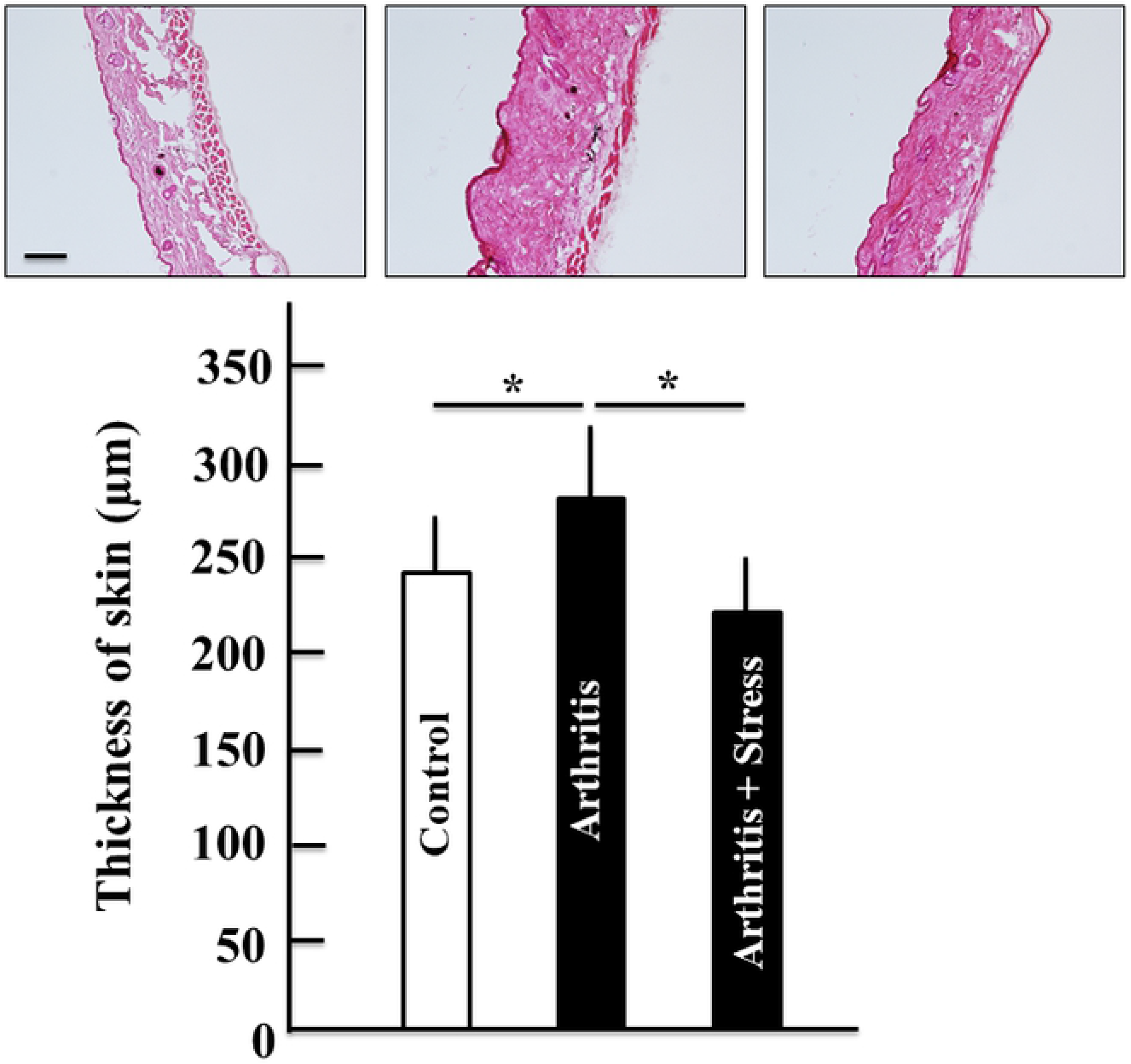
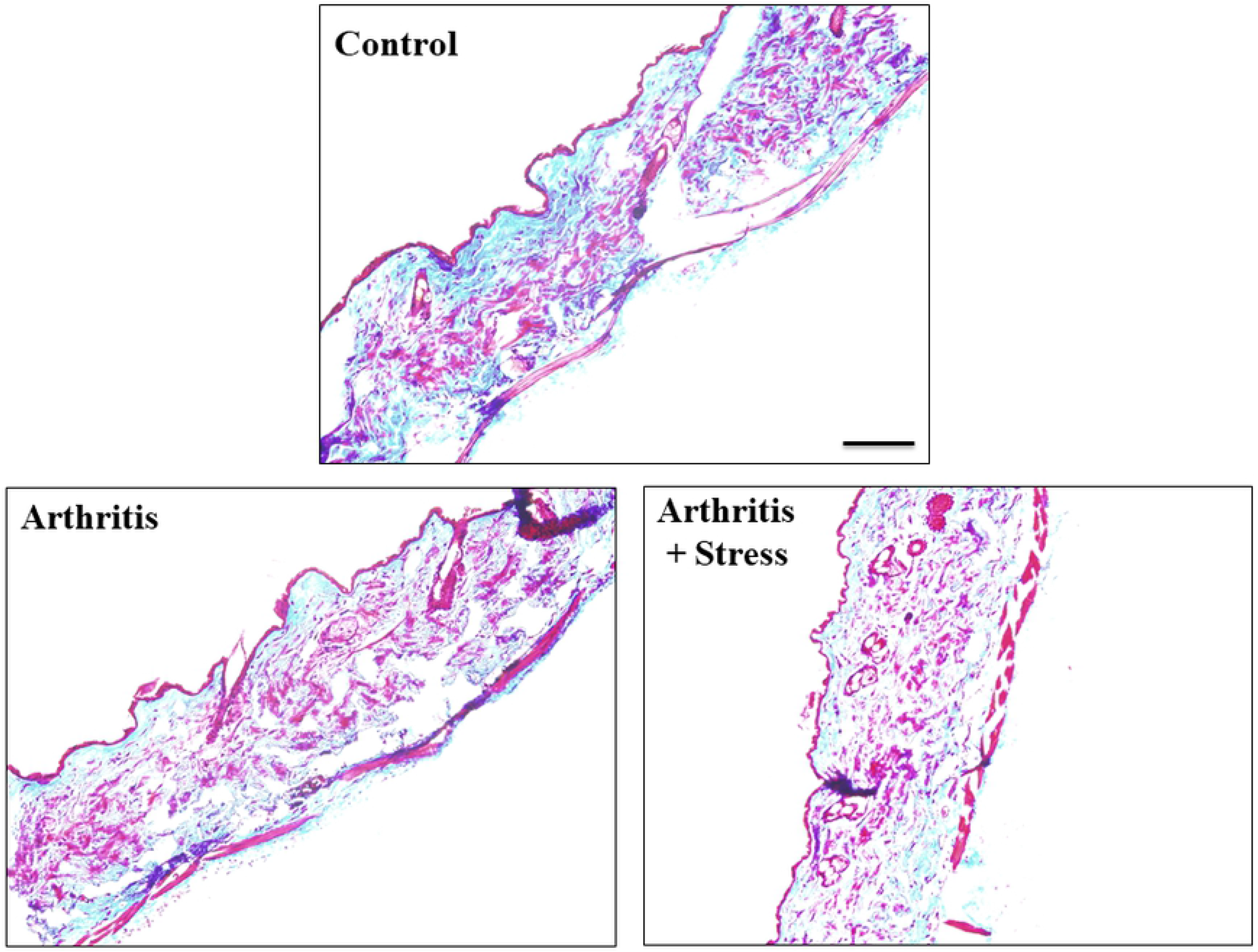
Changes in skin condition after application of stress in arthritis mice. A) Measurement of transepidermal water loss (TEWL) on the dorsal skin of mice on the final day. B) Hematoxylin and eosin (H&E) staining of the skin tissue section. Scale bar = 100 μm. Overall skin thickness was estimated based on the average length from the outer layer of the epidermis to the border of the subcutis in 10 randomly selected fields for each sample. C) Masson trichrome staining of the skin tissue section. Scale bar = 100 μm. Values represent the rate of increase compared with the control mice and are presented as means ± standard deviations. Data are presented as means ± standard deviations (n = 5). \**P* < 0.05 (Tukey’s post-hoc test).

### Mast cell numbers in the skin

Our previous report suggested that mast cell activation induced skin dryness [10]. Therefore, we assessed the number of mast cells in each group using histological staining with toluidine blue. Notably, the number of mast cells was increased in arthritic mice compared with that in control mice (Fig. 3), but that in stressed arthritic mice was not different compared with that in arthritic mice.

**Figure 3.**
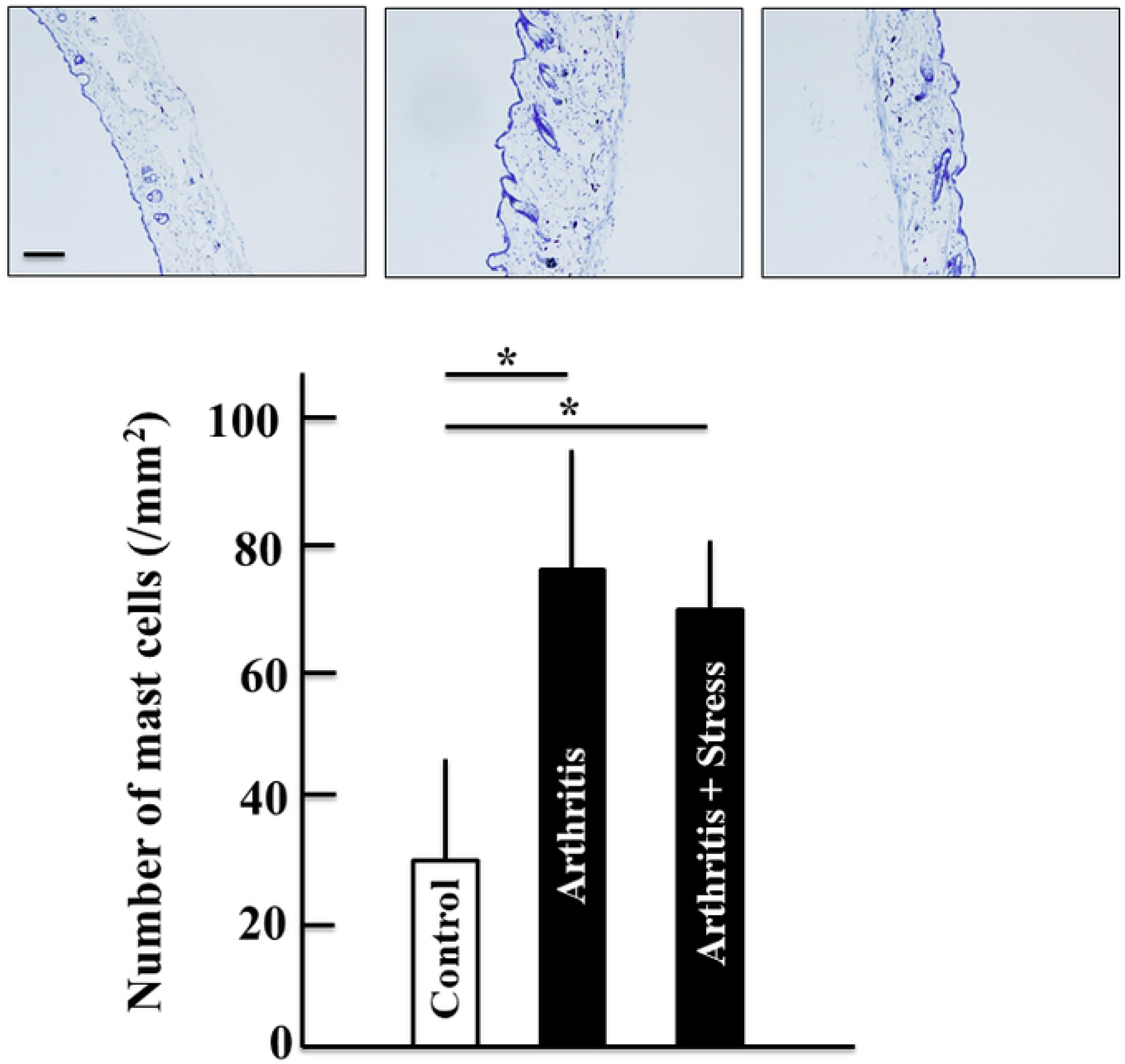
Effects of water immersion stress treatment on the number of mast cells in the skin of arthritis mice. The numbers of mast cells in arthritic mice and treated mice were counted following toluidine blue staining of skin specimens. The number of mast cells was counted in toluidine blue-stained skin tissue specimens. Scale bar = 100 μm. Data are presented as means ± standard deviations (n = 5). \**P* < 0.05 (Tukey’s post-hoc test).

### ROS, IL-6, corticosterone, and hyaluronan levels

Next, we examined the concentrations of ROS, IL-6, corticosterone (a stress hormone), and hyaluronan (a natural moisture factor) in the plasma (Fig. 4). The levels of ROS and corticosterone were increased in arthritic mice compared with those in control mice. Although ROS levels did not change in stressed arthritic mice, the levels of IL-6 and corticosterone were increased compared with those in arthritic mice. In contrast, hyaluronan levels were decreased in stressed arthritic mice compared with those in control mice. However, there were no differences in hyaluronan levels between arthritic mice and stressed arthritic mice.

**Figure 4.**
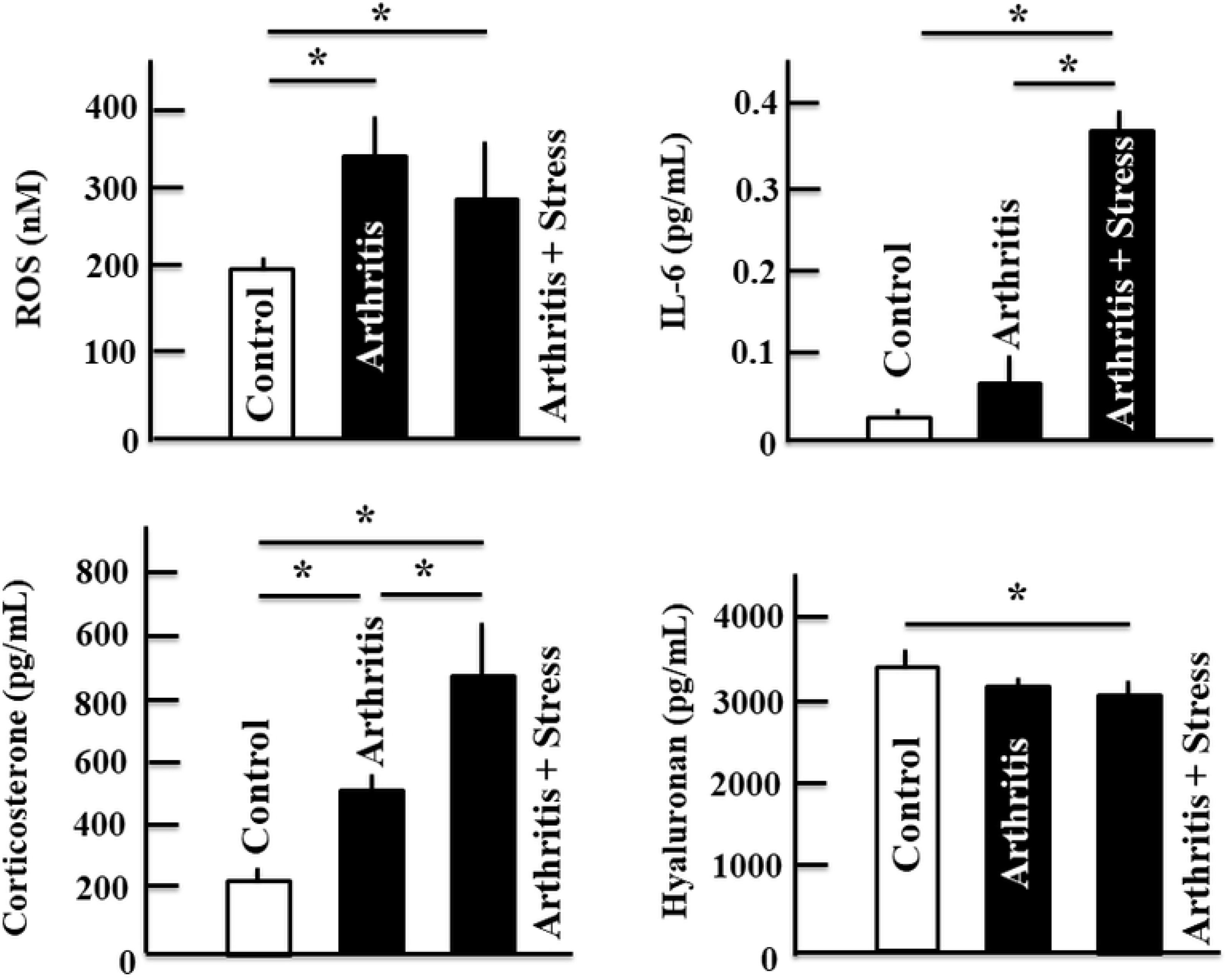
Effects of water immersion stress treatment on the plasma levels of ROS, IL-6, corticosterone, and hyaluronan in arthritic mice. Plasma levels of these proteins were assessed by enzyme-linked immunosorbent assay. Data are presented as means ± standard deviations (n = 5). \**P* < 0.05 (Tukey’s post-hoc test).

### Changes in thymus and spleen weights

Thymus weights in arthritic mice were decreased compared with those in control mice and significantly decreased in stressed arthritic mice (Fig. 5). In contrast, spleen weights were increased in arthritic mice compared with those in control mice and tended to increase in stressed arthritic mice compared with those in arthritic mice.

**Figure 5.**
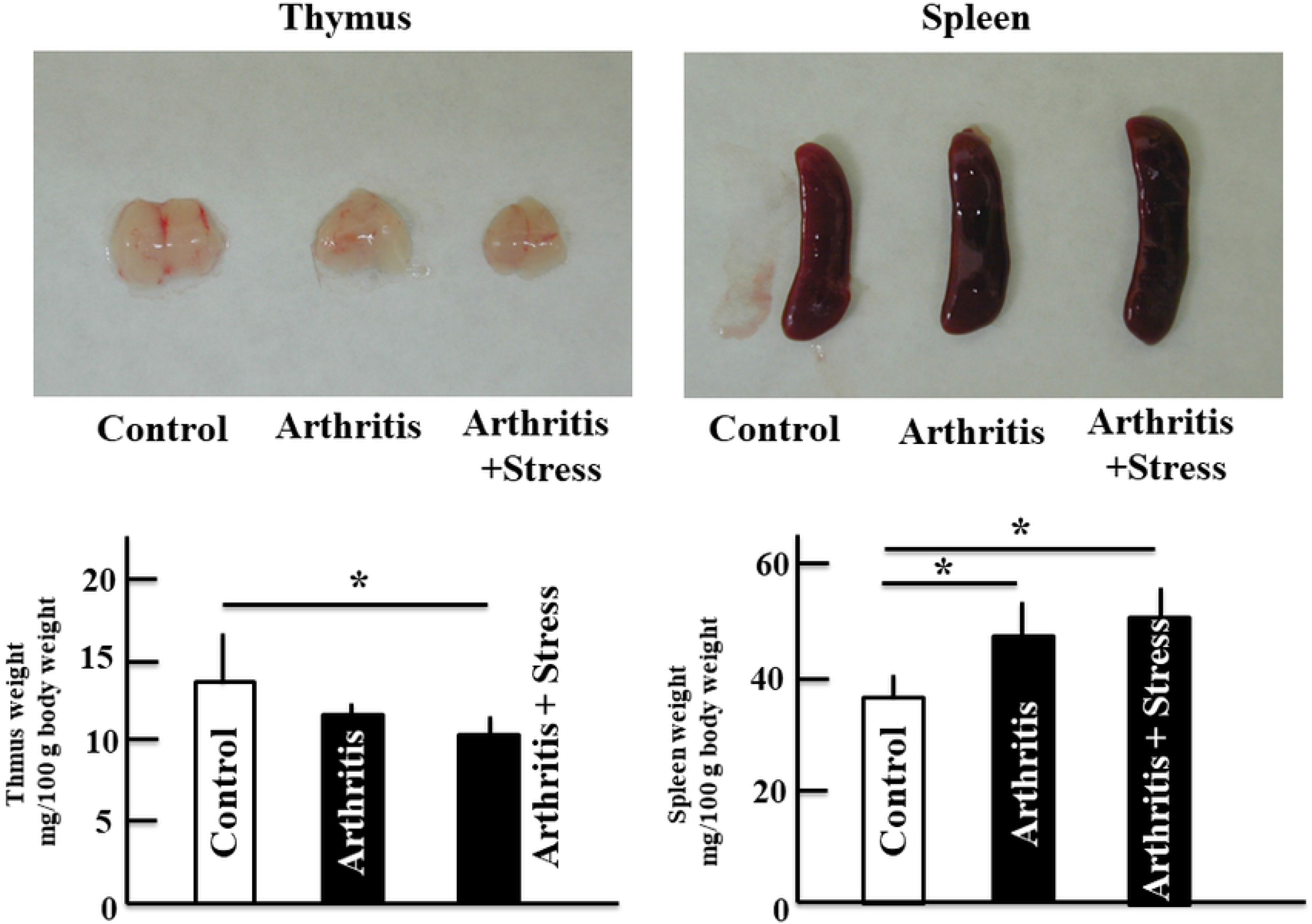
Effect of water immersion stress treatment on the weight of thymus and spleen in the arthritis mice. Data are presented as mean ± standard deviation (n = 5). *P < 0.05 (Tukey’s post hoc test).

**Effects of stress on Annexin V, glucocorticoid receptor (GCR), T-box protein expressed in T cells (T-bet), GATA binding protein 3 (GATA3), retinoic acid-related orphan receptor gamma t (RORγt), and Forkhead box protein 3 (Foxp3) in the thymus and spleen**

Annexin V detects changes in the cell membrane induced by apoptosis. Thymic annexin V expression was increased in arthritic mice compared with that in control mice and further increased in stressed arthritic mice. However, splenic annexin V expression did not differ among groups (Fig. 6B).

**Figure 6.**
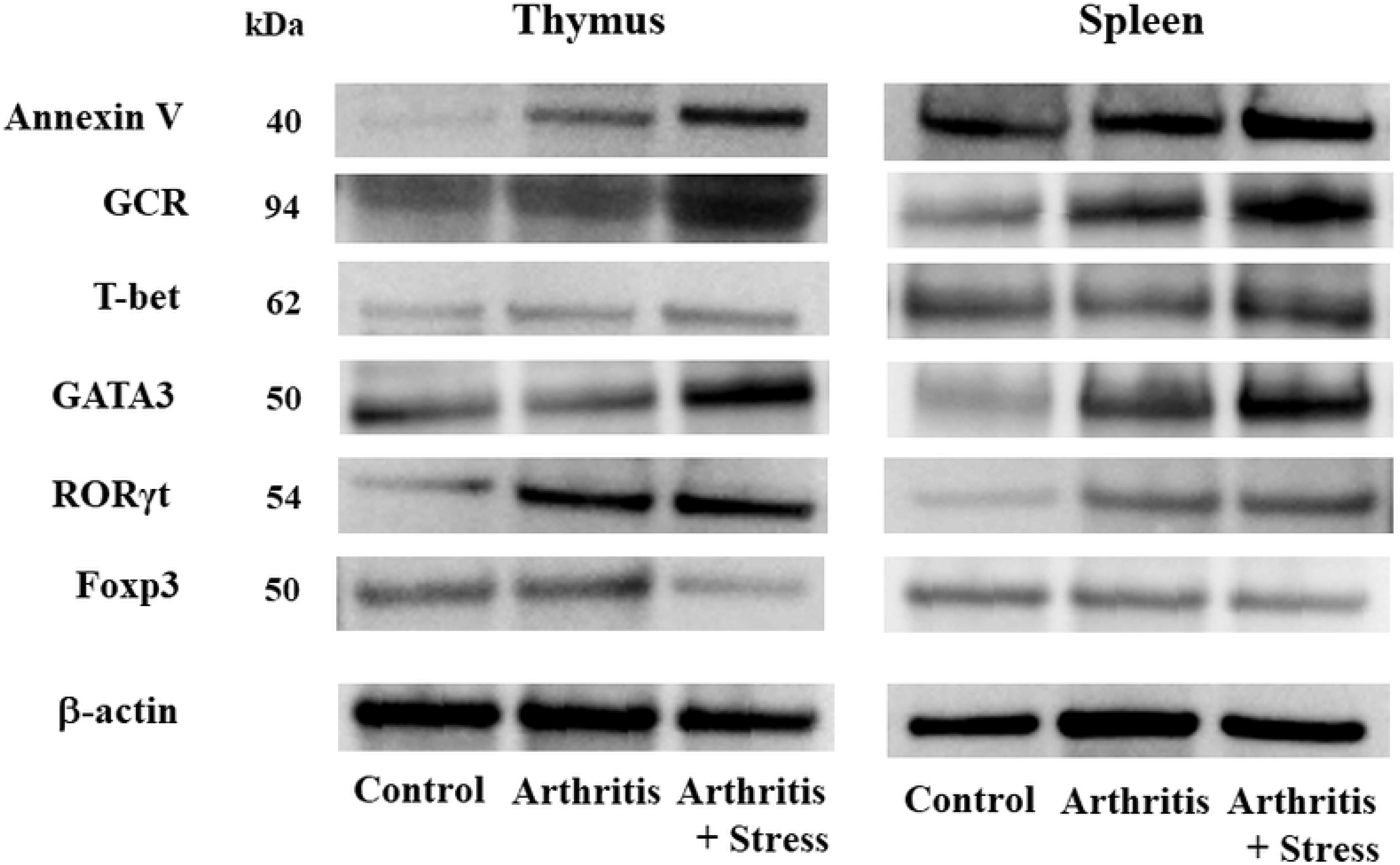
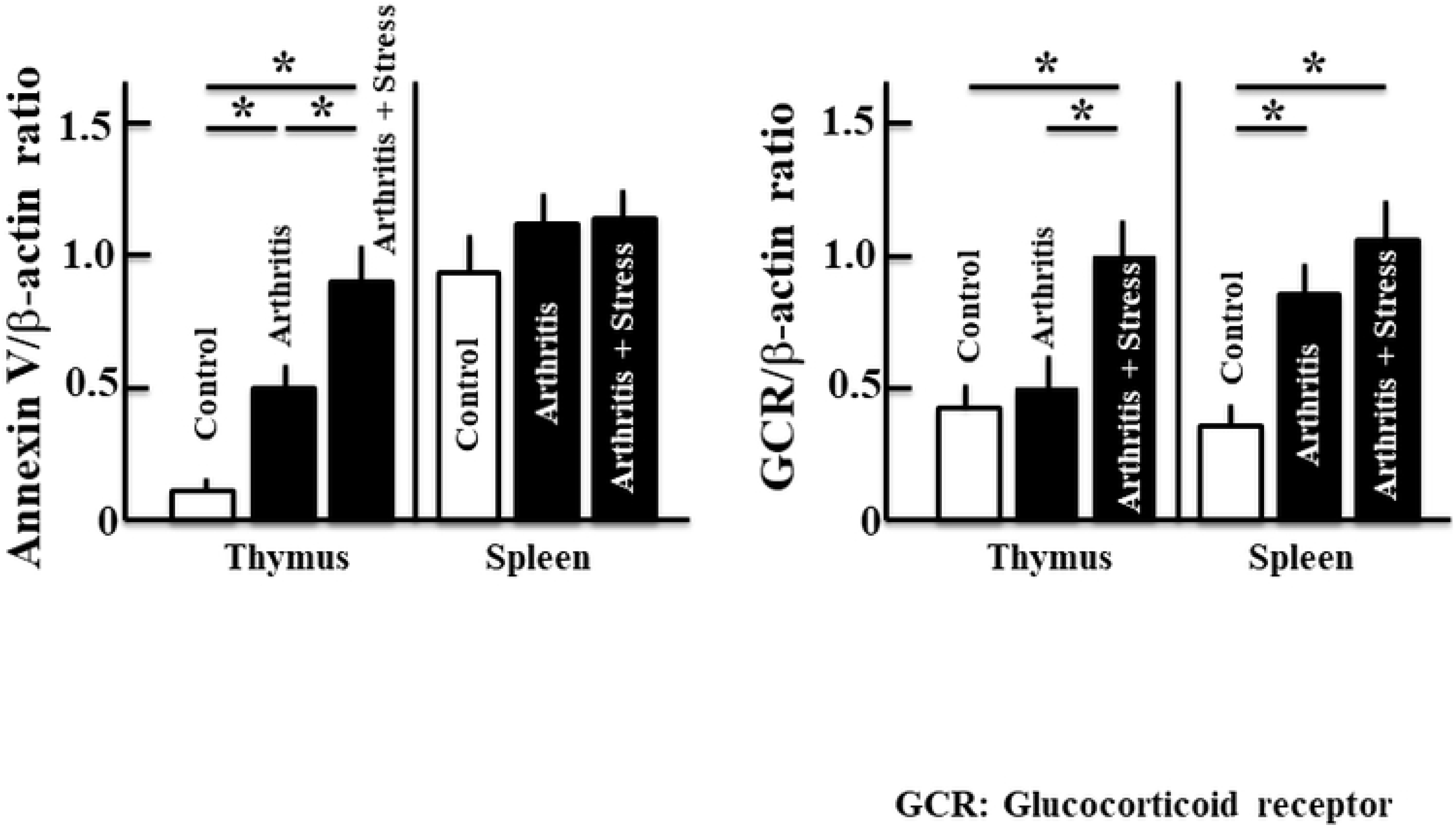
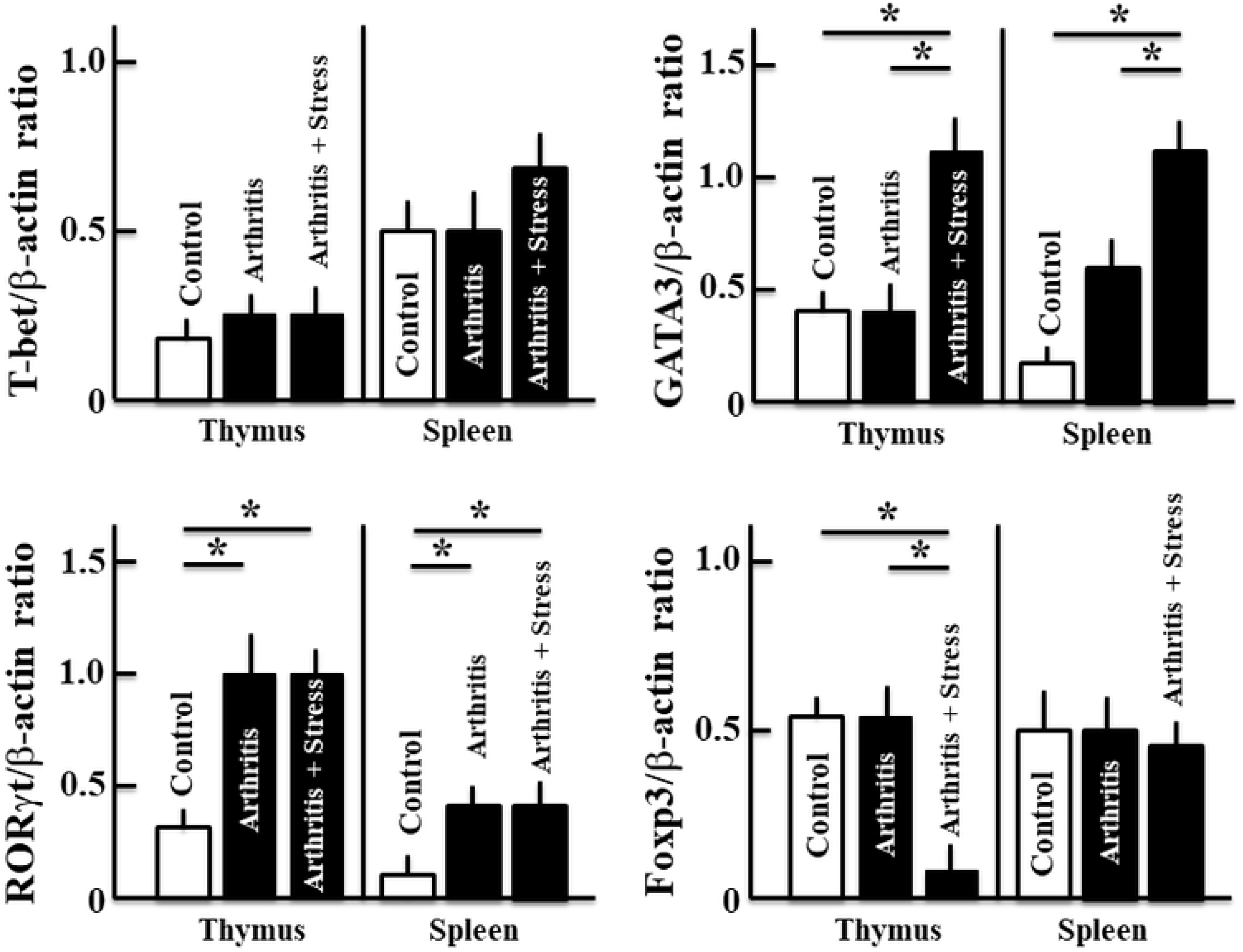
Effect of water immersion stress treatment on the expression of Annexin V, GCR, T-bet, GATA3, RORγt, and Foxp3 in the thymus and spleen of arthritic mice. Protein expression was detected by western blotting, with β-actin serving as a loading control. A) Protein band images. B) The expression of Annexin V and GCR in the thymus and spleen of arthritic mice. C) The expression of T-bet, GATA3, RORγt, and Foxp3 in the thymus and spleen of arthritic mice. Data are presented as means ± standard deviations (n = 5). \**P* < 0.05 (Tukey’s post-hoc test).

Thymic GCR expression did not differ between arthritic and control mice but increased in stressed arthritic mice. In addition, splenic GCR expression was increased in arthritic mice compared with that in control mice and tended to increase in stressed arthritic mice (Fig. 6B).

T-bet, GATA3, RORγt, and Foxp3 are specific transcription factors in T helper (Th) 1 cells, Th2 cells, Th17 cells, and regulatory T cells (Tregs), respectively [17–20]. In this study, we found that T-bet expression did not differ in any of the groups (Fig. 6C). In contrast, although thymic GATA3 expression did not differ between arthritic and control mice, GATA3 was upregulated in stressed arthritic mice compared with that in the other two groups. Splenic GATA3 expression was increased in arthritic mice compared with that in control mice and further increased in stressed arthritic mice (Fig. 6C). Additionally, thymic and splenic RORγt expression was increased in arthritic mice compared with that in control mice, but unchanged in stressed arthritic mice compared with that in arthritic mice (Fig. 6C). Finally, thymic Foxp3 expression did not differ between arthritic and control mice, but was decreased in stressed arthritic mice compared with that in the other groups. No differences in splenic Foxp3 expression were observed among groups (Fig. 6C).

## DISCUSSION

In this study, we evaluated the relationship between physical stress and induction of dry skin in arthritis using RA model mice. Physical stress load was applied by water immersion stress. In this model, we observed increased TEWL and decreased collagen compared with that in unstressed arthritic mice, indicating increased skin dryness. Additionally, plasma levels of ROS and corticosterone were increased in arthritic mice compared with those in control mice. Plasma levels of IL-6 and corticosterone were increased in stressed arthritic mice compared with those in unstressed arthritic mice. We also observed decreased Treg numbers and increased Th2 cell numbers in the thymus of stressed arthritic mice compared with that in unstressed arthritic mice.

We previously reported that various factors, including mast cells, ROS, collagen, and hyaluronan, were involved in skin dryness [10, 11, 21, 22]. Therefore, we evaluated the relationships of skin dryness with these factors. Importantly, we found that there were no changes in mast cell numbers, ROS levels, and hyaluronan levels among the different groups of mice. Thus, dry skin in stressed arthritic mice may not be related to these factors. However, the levels of IL-6, which exerts various effects on the skin [23, 24], were markedly increased in stressed arthritic mice. Therefore, IL-6 may be related to exacerbation of dry skin in the context of stress plus arthritis.

Notably, we found that skin dryness in arthritic mice was ameliorated by treatment with the GCR inhibitor RU486 [11]. This result suggested that GC is related to dry skin in arthritis. Consistent with this, secretion of GC is increased by various physical stresses [25, 26]. Therefore, in this study, we examined arthritis- and stress-induced skin dryness in mice using water immersion stress. The findings showed that stress worsened arthritis-induced dry skin. Moreover, we showed that the stress hormone GC (i.e., corticosterone) was increased in arthritic mice compared with that in control mice, and further increases were observed in stressed arthritic mice. Stress worsens skin diseases, such as psoriasis, and this phenomenon could be related to defects in the hypothalamic-pituitary-adrenal (HPA)-axis, altered glucocorticosteroid availability, or GCR dysfunction [27]. Because corticosterone generally has immunosuppressive effects, this hormone decreases IL-6 levels but stimulates; however, IL-2 plus corticosterone treatment increases IL-6 [28]. Thus, the specific relationship between IL-6 and corticosterone is unclear. In the current study, corticosterone and IL-6 levels were increased following application of stress in arthritic mice. Accordingly, these findings suggested that corticosterone and IL-6 were closely related and that IL-6 production may have been induced by corticosterone.

We then evaluated the levels of corticosterone and other factors in the thymus and spleen. The thymus is an organ involved in the proliferation and maturity of T cells as immune-competent cells. Stimulation of the HPA axis by stress results in corticosterone production, leading to thymus apoptosis and atrophy [29, 30]. Our current findings showed that increased corticosterone and GCR levels in the thymus may induce thymic apoptosis and significant thymic atrophy in stressed arthritic mice. The spleen is the largest organ in the lymphatic system and functions to remove old red blood cells and store immune cells, such as B and T cells [31]. In this study, the spleen size in arthritic mice was larger than that in control mice, and that in stressed arthritic was larger than that in arthritic mice. Splenomegaly is a symptom of Felty’s syndrome, which accompanies the autoimmune disease RA, resulting in disruption of skin health [32]. Restraint stress also decreases innate immune function, particularly natural killer cell activity and chemokine and cytokine expression in the spleen [33]. Therefore, immune cells and stress in autoimmune diseases are closely related to each other, and deterioration of dry skin in stressed arthritic mice may involve the thymus and spleen, although the details are unclear.

In this study, we observed the expression of Th2 cell- and Th17 cell-specific transcription factors related to the induction of dry skin in arthritic mice [21], and we confirmed the expression of Th1 cell- and Treg-specific transcription factors involved in Th2 and Th17 cells. T-bet, a Th1 cell-specific transcription factor, functions to provide protection against intracellular pathogens and viruses and produce interferon γ and IL-2 [17]; we found that T-Bet expression did not change in the thymus or spleen in any of the groups of mice. Therefore, Th1 cells were not related to dry skin in arthritic mice and were not also related to deterioration of dry skin in stressed arthritic mice.

RORγt is a transcription factor expressed in Th17 cells, which produce IL-17, IL-21, and IL-22 and differ functionally from Th1 and Th2 cells [34]. Our results showed that this protein was upregulated in the thymus and spleen in arthritic mice; however, no changes were observed in stressed arthritic mice. We have previously shown that mast cells are deeply involved in the induction of dry skin in arthritis through modulating the functions of Th17 cells [21]. However, our current results suggested that Th17 cells may not be related to deterioration of dry skin in stressed arthritic mice.

GATA3, a transcription factor in Th2 cells, is involved in allergic responses and can produce an array of cytokines, including tumor necrosis factor-α, IL-4, and IL-6, upon activation [18, 35]. Our results showed that GATA3 expression was increased in both the thymus and spleen in stressed arthritic mice compared with that in arthritic mice, suggesting that stress plus arthritic exacerbated dry skin in mice through modulation of Th2 cells. Therefore, we then evaluated the numbers of Tregs, which control T cells, in order to explore the cause of increased Th2 cell numbers in stressed arthritic mice. Tregs play a major role in the development of peripheral immune tolerance, and defects in Treg numbers lead to severe autoimmune diseases. These functions also have some detrimental effects on antitumor immunity [36]. Our results showed that the numbers of Tregs in the thymus were not altered in arthritic mice compared with that in control mice; however, a significant decrease was observed in stressed arthritic. Notably, no changes were observed in the spleen. The observed decrease in Tregs could disrupt the modulation of Th cells and result in abnormalities in the expression of inflammatory cytokines [37]. In the current study, thymus atrophy (apoptosis) was caused by increased corticosterone levels and was thought to decrease the expression of Tregs. Th cells were thought to lose this regulatory function owing to a reduction in Tregs; this would result in overexpression of Th2 cells and increased IL-6 levels, thereby. Increased IL-6 induces the degradation of collagen in the skin [38, 39]. Therefore, in stressed arthritic mice, this mechanism may have contributed to the deterioration of dry skin. In the spleen, Treg numbers were not altered between arthritic mice and stressed arthritic mice. However, Th2 cell numbers were increased in stressed arthritic mice. These results may be consistent with the increased expression of IL-6 in the spleen. Further studies are needed to elucidate the detailed mechanisms.

In summary, our results suggested that abnormalities in the immune system resulted in dry skin in stressed arthritic mice. Stresses vary from being acute to chronic, and can be physical or mental. In this study, we applied physical stress in mice, which is difficult to quantify, and hence was a limitation of this experiment. However, because patients with RA tend to experience both mental and physical stresses, it is necessary to reduce them in order to prevent skin diseases. Future studies are needed to examine the effects of reduced psychological stress on symptoms of RA, patient quality of life, and RA-related skin diseases.

## Acknowledgements

This study was supported by JSPS KAKENHI (Grant Number 18K06082)

